# Short-Term Performance Assay Identifies Functional Benefits and Early Toxicity of Longevity Interventions in Mice

**DOI:** 10.64898/2026.02.25.707674

**Authors:** Elisa Marín-Jerez, Javier Rueda-Carrasco, Florinda Meléndez-Rodríguez, Patricia Partido-Borge, Elena Tapia, Benjamin D. Leibowitz, Alberto Parras

**Author notes:** Corresponding autor. Co-first authors (Elisa Marín-Jerez, Javier Rueda-Carrasco).

## Abstract

Mouse lifespan studies are slow and costly, limiting the number of interventions that have demonstrated robust anti-aging effects. This highlights the need for rapid early-stage screening tools capable of assessing both efficacy and potential side effects.

Here, we present a short-term performance assay designed to rapidly profile functional benefits and early toxicity of longevity interventions in mice. Over an 8-week period, mice received one of five candidate anti-aging treatments: 17α-estradiol, rapamycin + Smer28, berberine + resveratrol, sildenafil and pinealon. The protocol longitudinally monitored body weight and temperature, and food intake, alongside post-treatment assessments of grip strength, locomotor activity, Y-maze cognition, social behavior, and hematological and urinary parameters.

The screen revealed compound-specific phenotypes: 17α-estradiol induced significant weight loss, increased grip strength, and dorsal alopecia, consistent with metabolic remodeling. Sildenafil reduced basal body temperature and preserved locomotor activity. Berberine + resveratrol decreased food intake and fasting glucose without major changes in physical performance, resembling caloric restriction-like metabolic effects. Rapamycin + Smer28 modestly improved strength and sociability but induced anemia in 2 of 5 mice, indicating potential dose-dependent toxicity. Pinealon showed a trend toward improved working memory without detectable adverse effects.

This multi-parametric approach enables discover healthspan extending interventions facilitating prioritization and dose refinement before committing to full lifespan studies.

Finally, to our knowledge, this represents the first comprehensive preclinical aging study in mice fully funded through tokenized decentralized science (DeSci), demonstrating how community-governed, on-chain funding can support resource-intensive *in vivo* research.

## INTRODUCTION

Aging is accompanied by progressive declines in physiological function and an increased risk of chronic disease. Caloric restriction (CR) remains the gold-standard longevity intervention, extending lifespan across organisms from yeast to mammals (Sun et al., 2009).

In the last years, a rapidly growing number of anti-aging candidate interventions are being proposed through *in silico* approaches, including artificial intelligence-based pipelines, as well as through screening *in vitro* and in short-lived organisms such as yeast, worms, flies, or killifish (Bene & Salmon, 2023; Kim et al., 2016). However, translation of these candidates to mammals remains limited, as robust lifespan validation in rodents are costly (exceeding $200K USD), time-consuming (lasting over 2 years), and ethically demanding due to the large cohorts required (∼100 animals) (Harrison et al., 2019, 2021; Miller et al., 2011; Xu et al., 2021; Zhavoronkov et al., 2025).

These logistical demands have not only limited the number of compounds evaluated but also restricted the mechanistic pathways that can be rigorously tested, creating a major translational bottleneck in aging research (Malavolta, 2025; Marcozzi et al., 2025; Mitchell et al., 2016). Particularly in mice, it has been demonstrated that both pharmacological and genetic interventions can extend lifespan and improve healthspan (Harrison et al., 2019, 2021; Miller et al., 2011; Xu et al., 2021). For example, several conserved pathways implicated in aging can be targeted pharmacologically, including mTOR signaling, nutrient sensing, mitochondrial function, and vascular homeostasis (Grunewald et al., 2021; Mercken et al., 2014; Miller et al., 2011; Mills et al., 2016). Rapamycin remains a benchmark geroprotector, consistently extending lifespan across multiple strains and sexes (Harrison et al., 2009; Strong et al., 2020; Wilkinson et al., 2012). However, 17α-estradiol robustly extends lifespan in male mice, but not in females, and is associated with pronounced reductions in adiposity and sex-specific metabolic responses (Harrison et al., 2014, 2021; Strong et al., 2016). Additional interventions, including glucose-lowering agents (acarbose, metformin, berberine, or canagliflozin) and senescence-modulating strategies, have also been shown to improve metabolic health, frailty-related outcomes, and survival in mice (Dang et al., 2020; Harrison et al., 2014, 2019; Martin-Montalvo et al., 2013; Miller et al., 2020). Notably, combinations of interventions, such as rapamycin with metformin or acarbose, can synergistically enhance lifespan benefits, highlighting the importance of systematically evaluating multi-compound regimens (Strong et al., 2016, 2022). Finally, some promising interventions that appear beneficial may induce secondary toxicity with chronic use, underscoring the need to detect adverse effects early during preclinical evaluation (Harrison et al., 2024; Palliyaguru et al., 2020; Strong et al., 2022).

Although recent efforts have proposed standardized protocols for conducting scalable mouse longevity studies (Zhavoronkov et al., 2025) and more ethically refined strategies based on phenotyping in standard home-cage have been developed (Delalić et al., 2025), a significant gap between rapid intervention discovery and definitive validation in mouse lifespan studies still remains.

To address this gap, we developed a short-term, performance-based assay in middle-aged male C57BL/6J mice to rapidly profile pharmacological interventions for functional benefits and early toxicity signals. Over an 8-week treatment period, we longitudinally quantified body weight, core temperature and food intake, and assessed multiple functional domains including grip strength, locomotor activity, cognition, and social behavior, alongside hematological and biochemical measures. As proof-of-concept, we tested five interventions linked to aging-associated pathways: 17α-estradiol; rapamycin combined with the autophagy enhancer Smer28; berberine plus resveratrol; sildenafil; and pinealon. By combining multiple health-related functional measures with basic clinical safety assessments within a short chronic timeframe, this protocol helps prioritize promising anti-aging interventions, identify targeted pathways, and detect potential side effects or dosing limitations before committing to full mouse lifespan studies.

## RESULTS

### Longitudinal monitoring of basic physiological parameters

Seven-month-old male C57BL/6J mice were acclimated for four weeks prior to being randomly assigned to six experimental groups (n = 5 mice): Control (CTR), 17α-estradiol (EST; 22 ppm in food), berberine + resveratrol (B+RE; 375 ppm + 170 ppm in food), rapamycin + SMER28 (R+SM; 22 ppm + 88 ppm in food), sildenafil (SIL; 62 ppm in food), and pinealon (PNL; 30 mg/kg intraperitoneal biweekly) (Fig. 1a). During the acclimation period, one mouse from the sildenafil group required euthanasia due to fighting-related injuries, and the group was maintained with n = 4 thereafter. All remaining animals entered the longitudinal phenotyping protocol, which included an 8-week baseline period followed by 8 weeks of chronic treatment (Fig. 1b).

**Figure 1.**
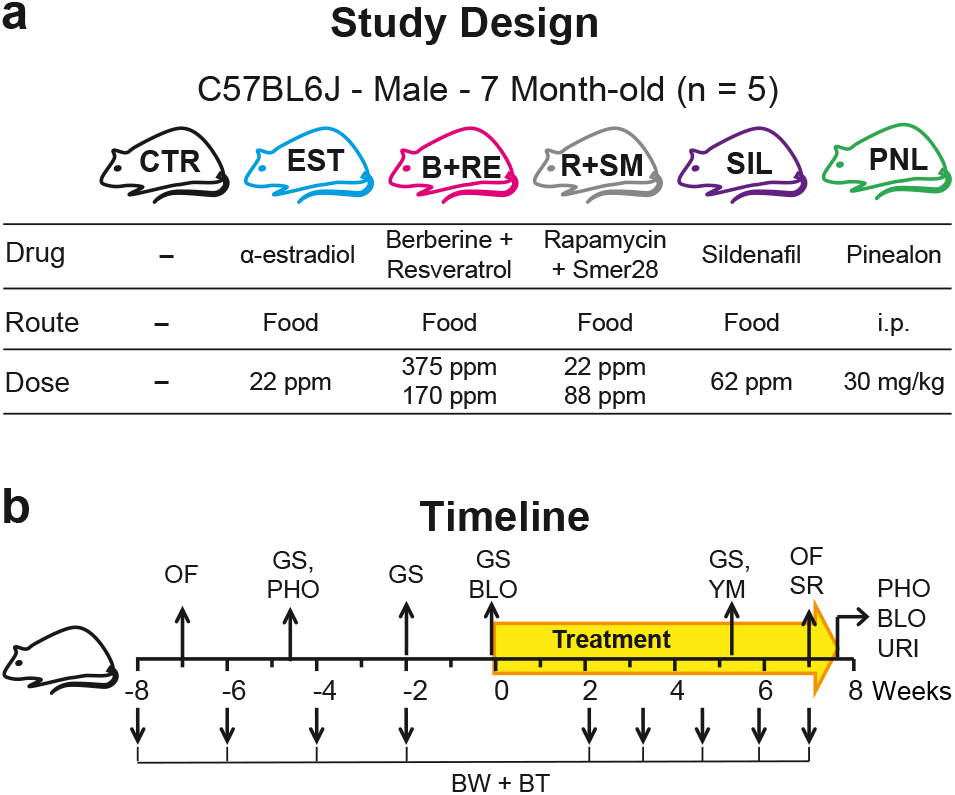
Experimental protocol. **a**, Study design showing mouse strain, treatment groups, compounds, administration route and dose for control and intervention groups (n = 5 per group, 7-month-old male C57BL/6J mice). **b**, Timeline of the 8-week protocol including behavioral tests activity by open field (OF), grip strength (GS), working memory by Y-maze (YM), social recognition (SR), physiological monitoring, body weight (BW), body temperature (BT), photographs (PHO), blood (BLO) and urine (URI) collection.

Body weight and temperature, and food intake were monitored as mandatory longitudinal parameters due to their relevance in aging research. Body weight and food intake reflect systemic metabolic adaptations and caloric restriction-like effects, while body temperature has been associated with lifespan regulation and aging trajectories (Acosta-Rodríguez et al., 2022; Conti & Cabo, 2025; Francesco et al., 2024). After treatment onset, most groups produced body weight curves comparable to controls (Fig. 2a and Extended Data Fig. 1a), however, as previously reported (Harrison et al., 2014, 2021; Strong et al., 2016), 17α-estradiol induced a rapid and significant body weight loss, consistent with its known effects on fat mass reduction (Bubak et al., 2024).

**Figure 2.**
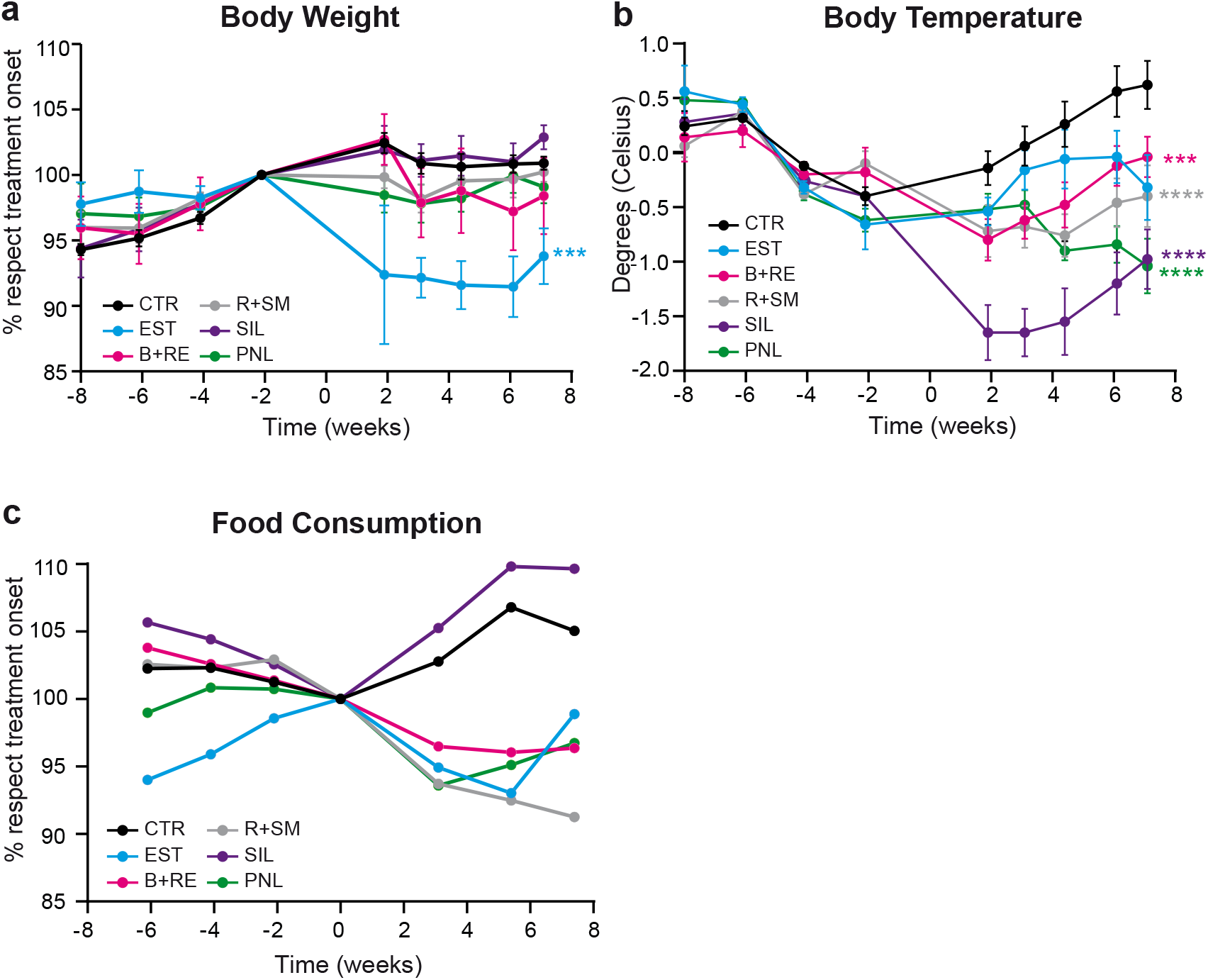
Basic physiological parameters. **a**, Body weight variation over time, expressed as percentage, **b**, Rectal body temperature variation over time, expressed as degrees Celsius (ºC) change relative and **c**, Food consumption over time, expressed as percentage relative to treatment onset. CTR, control; EST, 17α-estradiol; B+RE, berberine + resveratrol; R+SM, rapamycin + Smer28; SIL, sildenafil; PNL, pinealon. Data show mean ± standard error of the mean (SEM). Statistical significance was assessed by two-way ANOVA with Dunnett’s multiple comparisons test (**a**) and Mixed-effects model (REML) with Dunnett’s multiple comparisons (**b**).

At baseline, all groups displayed comparable core body temperature values (∼36.0°C) (Extended Data Fig. 1b). Following treatment onset, sildenafil-treated mice exhibited a marked decrease in basal body temperature of more than 1°C, which persisted throughout the treatment period (Fig. 2b). This effect is consistent with sildenafil-mediated vasodilation, which may increase peripheral heat dissipation. Although no other intervention induced a comparably large reduction in basal temperature, the remaining treated groups displayed a mild but significant lower body temperature compared to controls (Fig. 2b). This result supports the hypothesis that even a modest decrease in core body temperature may represent a longevity-associated physiological signature (Conti & Cabo, 2025; Zhao et al., 2022).

On the other hand, food intake progressively increased in control mice over time, consistent with their gradual weight gain (Extended Data Fig. 1c). In contrast, treated groups exhibited distinct feeding trajectories after intervention onset (Fig. 2c). Sildenafil-treated mice increased their food consumption and remained above control levels throughout the whole experiment, likely to reflect higher energetic demands associated with drug-induced heat loss and compensatory thermogenesis. In contrast, the remaining group showed reduced food consumption relative to controls. In the estradiol-treated animals, this reduction was consistent with the observed weight loss, while in the other groups it may reflect reduced metabolic demand and/or mild calorie restriction-like effects induced by the interventions (Fig. 2c and Extended Data Fig. 1c).

Together, these longitudinal basic physiological measurements revealed distinct compound-specific metabolic profiles, providing an essential baseline framework to interpret the functional, behavioral, and biomolecular outcomes observed later in the study.

### Physical Performance: Strength and Activity

To evaluate physical functions, known to decline with aging (Shoji et al., 2016; K. Xie & Ehninger, 2023), we measured forelimb grip strength and open-field locomotor activity before and after the treatment period. At baseline, all groups displayed comparable grip strength values normalized to body weight which increased over time, consistent with normal maturation, as mice continue to gain muscle strength into middle age (Extended Data Fig. 2a). Notably, 17α-estradiol-treated mice exhibited a highly significant improvement in strength (Fig. 3a). This effect is likely explained by the reduction in adipose mass, resulting in increased strength relative to body weight and suggesting that lean mass was preserved. The rapamycin + Smer28 group also showed a modest improvement in strength, whereas none of the other treatments produced significant changes compared to controls (Fig. 3a). Locomotor activity was assessed using the open-field test. At baseline, all mice displayed strong exploration behavior, with similar total distance traveled (∼30 m in 10 min) and comparable maximum speed across groups (Extended Data Fig. 2b-c). After 8 weeks, control mice exhibited a clear reduction in locomotor activity, a pattern also observed in most treatment groups. In contrast, the sildenafil group maintained high locomotor performance, both distance traveled and maximum speed were preserved and even increased compared to baseline (Extended Data Fig. 2b-c), resulting in significant improvement relative to all other groups (Fig. 3b). These findings suggest that sildenafil treatment preserved physical activity levels, which may be explained by increased blood flow and improved muscle perfusion induced by chronic vasodilation, potentially reducing fatigue and enhancing endurance.

**Figure 3.**
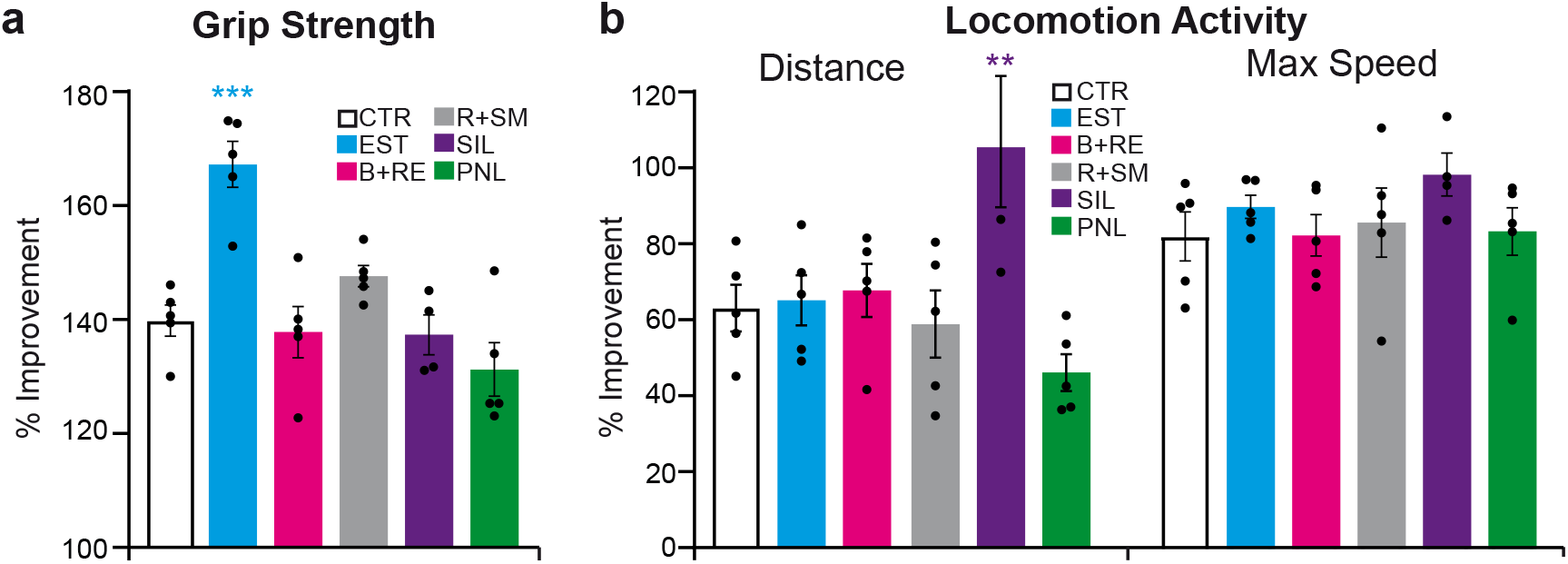
Physical Performance: Muscle strength and locomotor activity. **a**, Forelimb grip strength normalized by body weight and **b**, Locomotor performance in the open field test expressed including total distance traveled and maximum speed as percentage improvement relative to baseline. CTR, control; EST, 17 α-estradiol; B+RE, berberine + resveratrol; R+SM, rapamycin + Smer28; SIL, sildenafil; PNL, pinealon. Data show mean ± standard error of the mean (SEM). Statistical significance was assessed by one-way ANOVA with Dunnett’s multiple comparisons test **(a–b)**.

Together, these results indicate that sildenafil and 17α-estradiol, and to a lesser extent rapamycin + Smer28, promoted sustained or enhanced physical performance in middle-aged mice, supporting their potential relevance for healthspan-related outcomes.

### Cognitive Function and Social Behavior

Next, we assessed cognitive performance using the Y-maze spontaneous alternation test, that evaluates spatial working memory based on the natural tendency of mice to explore a novel arm rather than returning to a previously visited one. No significant differences were observed between control and treated groups in the total number of arm entries or in the percentage of spontaneous alternations (Fig. 4a-b), indicating preserved working memory across all conditions. Only pinealon-treated mice showed a trend toward increased alternation performance, consistent with its proposed nootropic properties (Khavinson et al., 2021); however, given the limited sample size and the relatively young age of the animals, this effect remained non-significant and may require longer treatment duration or larger cohort size to be confirmed (Fig. 4b).

**Figure 4.**
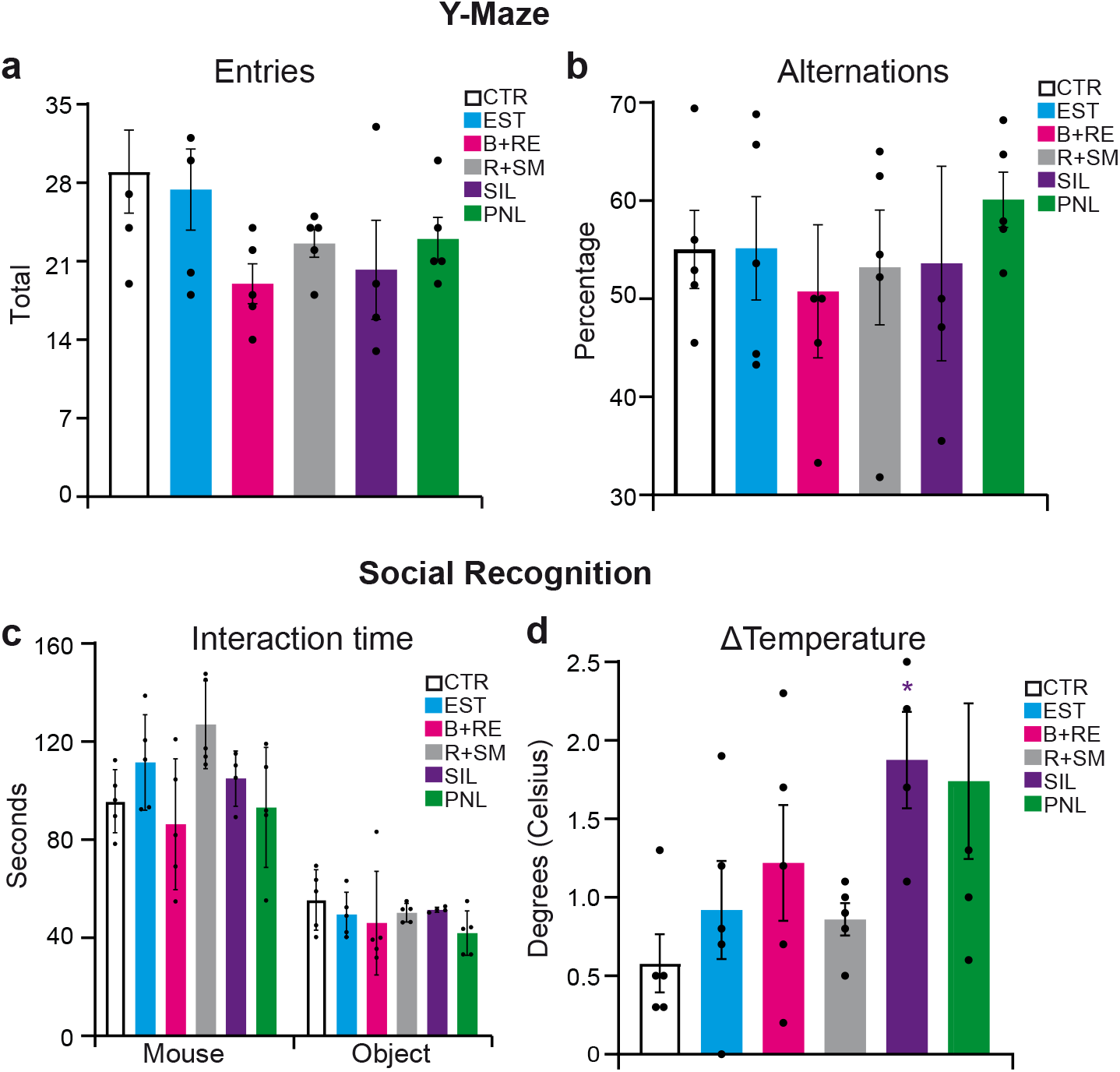
Cognition function and Social Behavior. **a**, Total number of arm entries and **b**, Percentage of spontaneous alternations in the Y-maze test. **c**, Time spent interacting with unfamiliar mice or inanimate objects during the 7 min of the social recognition test. **d**, Change in core body temperature (ΔT) measured before and after the social recognition test. CTR, control; EST, 17α-estradiol; B+RE, berberine + resveratrol; R+SM, rapamycin + Smer28; SIL, sildenafil; PNL, pinealon. Data show mean ± standard error of the mean (SEM). Statistical significance was assessed by one-way ANOVA with Dunnett’s multiple comparisons test (**a**,**b**,**c-mouse**) and Kruskal-Wallis test with Dunn’s multiple comparisons test (**c-object**,**d**).

Large population-based surveys show that with advancing age, rates of sexual activity fall and problems with sexual interest and arousal become increasingly common in both men and women (Lindau et al., 2007; Wang et al., 2015), and social interaction has been reported to decline in middle-aged mice (Boyer et al., 2019). Sexual and social behavior were evaluated by measuring the total time spent actively investigating two unfamiliar mice (one male and one female) compared to two inanimate objects during a 7-min trial. As expected, control and treated mice spent more time interacting with stranger mice than with objects (Fig. 4c and Extended Data Fig. 3a); however, the rapamycin + Smer28 group displayed increased social investigation time (Fig. 4c). Although no significant preference for male versus female conspecifics was detected in any group, including controls; rapamycin + Smer28 mice showed a trend toward increased interaction with females (Extended Data Fig. 3c-d), suggesting increased socialization with a potential preference toward females, which may reflect enhanced socio-sexual motivation.

During the social interaction test, we made an interesting observation regarding body temperature dynamics. Core body temperature was measured immediately before and immediately after the social recognition test to calculate the temperature change (ΔT) induced by social interaction and locomotor activity. All groups showed an increase in body temperature during the test (Extended Data Fig. 3b). Notably, the sildenafil group, which exhibited a significantly lower basal body temperature at rest (Fig. 2b), showed the largest increase in core temperature after the social test activity (Fig. 4d). This thermogenic response suggests that the reduced resting body temperature is not due to impaired heat production capacity, but rather to increased heat dissipation, consistent with the vasodilatory properties of sildenafil. These findings further support that the reduced basal temperature in this group did not compromise overall vitality or the ability to mount a physiological response during activity.

Overall, within this 2-month timeframe, no clear cognitive impairment was detected across treatments. Pinealon-treated mice displayed a trend toward improved working memory, and rapamycin + Smer28 mice exhibited increased sociability with a trend toward female-directed interaction. Unexpectedly, sildenafil-treated mice exhibited a strong increase in core body temperature during behavioral activity, demonstrating preserved thermogenic capacity despite their reduced basal temperature.

### Blood and metabolic endpoint analysis

Before treatment and after 8 weeks of the onset mice were photographed, and blood and urine samples were collected prior to euthanasia. Complete blood count (CBC) analysis revealed that most treatment groups displayed hematological profiles comparable to controls, with no clear evidence of improvement or impairment relative to baseline, only a small increase in red blood cell parameters was observed in the berberine + resveratrol group (Fig 5a). However, when analyzing individual animals in greater detail, two mice in the rapamycin + Smer28 group exhibited abnormally low red blood cell (RBC) count, hematocrit, and hemoglobin levels (Supplementary Table 1). These two out of five animals met clinical criteria consistent with anemia, also supported by elevated red cell distribution width (RDW), a parameter commonly associated with this disorder and reflecting increased heterogeneity in red blood cell size. Although rapamycin alone has been reported to increase RBC count and hematocrit (Acar et al., 2020), the anemia observed in the combination with Smer28-treated mice highlights a potential safety concern at the dose used in this study. This result supports the relevance of short-term screening protocols to identify safe and effective dosing prior to initiating long-term lifespan studies.

**Figure 5.**
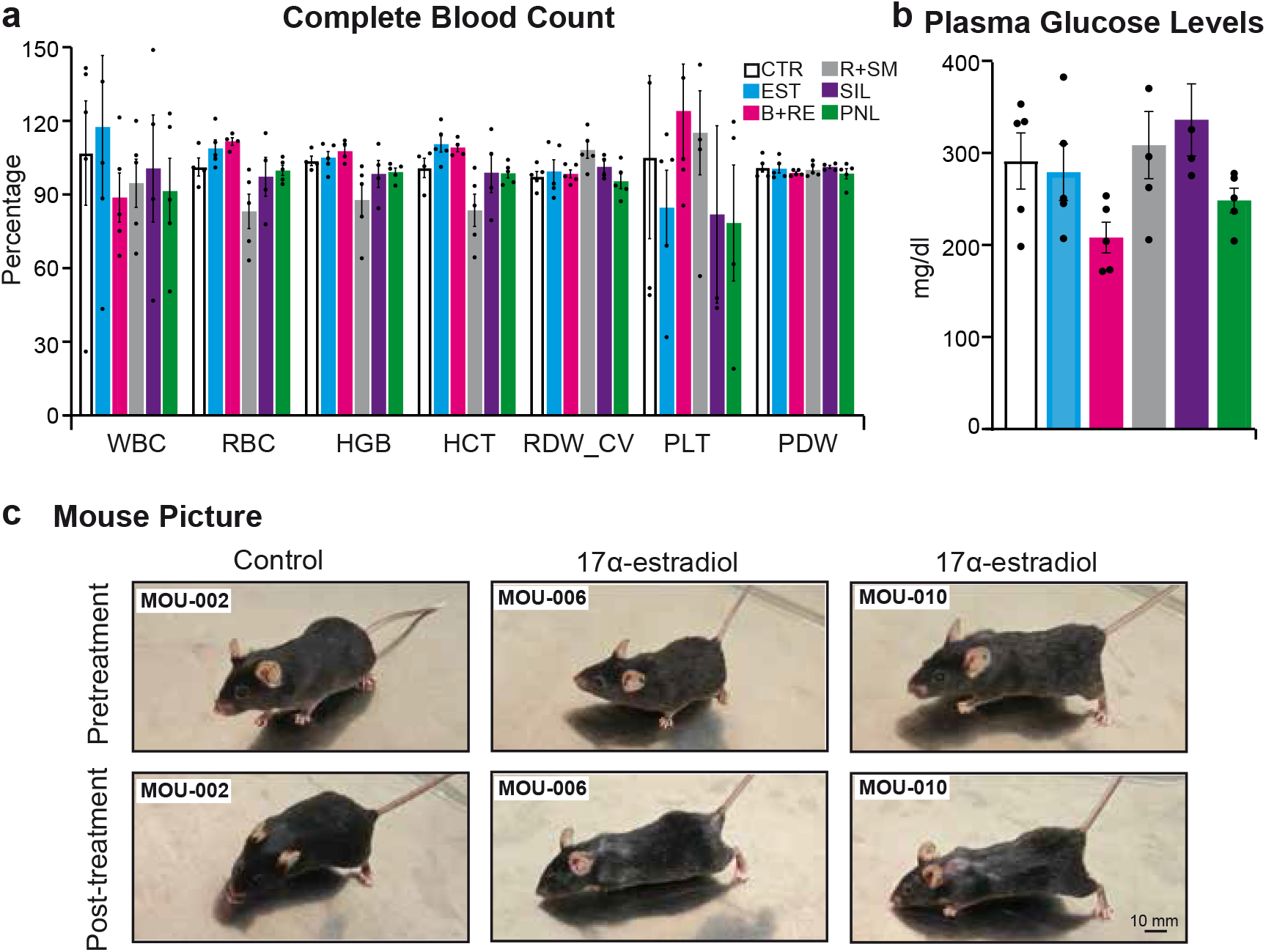
Blood and metabolic endpoint analysis. **a**, Complete blood count (CBC) parameters expressed as percentage relative to baseline. **b**, Fasting plasma glucose levels at study endpoint normalized by blood levels in the baseline. **c**, Representative photographs of mice before and after treatment, showing dorsal alopecia in the 17α-estradiol vs control group. CTR, control; EST, 17α-estradiol; B+RE, berberine + resveratrol; R+SM, rapamy-cin + Smer28; SIL, sildenafil; PNL, pinealon. Data show mean ± standard error of the mean (SEM). Statistical significance was assessed by one-way ANOVA with Dunnett’s multiple comparisons test **(a-b)**.

At the end of the study, fasting plasma glucose levels were lower in the berberine + resveratrol group (Fig. 5b), consistent with the well-described glucose-lowering effects of berberine (W. Xie et al., 2022). This finding highlights this combination not only as a potential longevity intervention, but also as a promising candidate for metabolic and diabetic disorders. No other treatment group showed significant differences in fasting glucose compared to controls (Fig 5b). In addition, a basic urinalysis was also performed at the endpoint using pooled urine samples from each treatment group. No abnormalities or relevant differences were detected across groups (Supplementary Table 2), suggesting the absence of detectable renal or urinary toxicity under the conditions tested.

Finally, comparisons of photographs taken before and after treatment revealed a clear alopecia phenotype in the 17α-estradiol group, predominantly affecting the dorsal region near the neck (Fig. 5c). This effect may be linked to estradiol-induced metabolic remodeling and potential activation of brown adipose tissue in this anatomical region; however, the mechanistic basis of this phenotype would require further investigation.

Taken together, this short-term longitudinal protocol identified distinct compound-specific physiological and functional profiles within 8 weeks of treatment. 17α-estradiol induced marked weight loss and alopecia, while sildenafil reduced basal core temperature but preserved locomotor activity and showed a strong activity-induced thermogenic response. Berberine + resveratrol showed a trend toward lower fasting plasma glucose, and pinealon showed a trend toward improved working memory. In contrast, rapamycin + Smer28 improved strength and sociability but induced anemia in two animals, highlighting a potential safety concern at the tested dose. Overall, these results support the utility of a cost-efficient short-term screening to identify early efficacy signals and detect adverse effects prior to long-term lifespan studies.

## DISCUSSION

The results obtained from this 4-month performance protocol in middle-aged mice indicate that an 8-week treatment window would be sufficient to reveal compound-specific patterns in physiology, functional capacity, and early toxicity. Rather than replacing lifespan trials, this approach serves as an initial screening step to identify interventions with favorable risk-benefit profiles before investing substantial resources in large-scale lifespan studies (Nadon et al., 2017; Zhavoronkov et al., 2025).

Using this framework, we observed distinct phenotypic signatures across the tested interventions, enabling a compound-specific evaluation of functional benefits and early safety liabilities. The 17α-estradiol group illustrates how established geroprotective patterns can be detected within a compressed timeframe. Previous work has consistently shown lifespan extension in male mice, accompanied by reductions in adiposity and sex-specific metabolic effects, with minimal benefit in females (Harrison et al., 2014, 2021; Strong et al., 2016). In our study, 17α-estradiol–treated mice exhibited rapid weight loss and increased grip strength normalized to body weight, consistent with a shift toward a leaner, functionally robust phenotype. The detection of alopecia as an overt phenotype further highlights the assay’s ability to capture potentially undesirable effects, informing dose and schedule considerations for future studies.

The berberine plus resveratrol combination produced a less pronounced phenotype, primarily characterized by metabolic changes. Berberine has been reported to ameliorate cellular senescence markers and extend lifespan in naturally aged mice (Dang et al., 2020), while resveratrol has been proposed as a caloric restriction mimetic with potential anti-aging properties, although its effects on lifespan in rodents remain controversial and context-dependent (Baur et al., 2006; Miller et al., 2011;

Pearson et al., 2008). In this short-term assay, the combination reduced food intake and lowered body temperature, in good agreement with the observed decrease in fasting plasma glucose, suggesting improved metabolic efficiency or a mild calorie restriction–like state. These findings illustrate how short-term assays can reveal beneficial metabolic adaptations that may precede detectable performance gains and highlight the importance of integrating biochemical with functional endpoints when prioritizing candidates for long-term testing (Mitchell et al., 2016; Xu et al., 2021).

Perhaps the most unexpected finding emerged from sildenafil treatment. Although PDE5 inhibitors such as sildenafil, are not considered canonical geroprotectors, vascular and perfusion pathways are increasingly recognized as key modulators of healthy aging. For example, VEGF-targeted strategies improve multi-organ function and survival in mouse models (Grunewald et al., 2021). Interestingly, two independent studies have reported that sildenafil use is associated with a reduced risk of Alzheimer’s disease (Fang et al., 2021) and with increased lifespan (Morin et al., 2024) in humans, supporting interest in PDE5 inhibition as a potential aging-relevant pathway. In our study, sildenafil-treated mice showed a robust preservation of locomotor activity when the rest of the groups exhibited a clear decline. This phenotype was accompanied by a marked reduction in basal body temperature clearly associated with promoting longevity (Conti & Cabo, 2025), while thermogenic responsiveness remained intact, as evidenced by a strong temperature increase during the social interaction test. One possible interpretation is that chronic vasodilation improves muscle perfusion, and reducing fatigue, thereby promoting a more “youth-like” activity profile despite lower resting temperature. Although the long-term lifespan effects of sildenafil remain unknown, these findings support further investigation of PDE5 inhibition as a potential healthspan-promoting strategy. It also illustrates how performance-based screening can uncover unexpected functional benefits from drugs originally developed for other clinical indications (Marcozzi et al., 2025; Mitchell et al., 2016).

In contrast, the rapamycin plue Smer28 combination highlights the critical role of early toxicity detection or dose adjustment. Rapamycin is the most consistently validated lifespan-extending drugs across organism, yet its benefits are sensitive to dose, schedule, and sex, and it is associated with metabolic side effects at higher exposures (Roark & Iffland, 2025). The addition of Smer28 aimed to potentially synergize with rapamycin in enhancing autophagy and proteostatic maintenance (Dang et al., 2020; Xu et al., 2021). However, at the chosen doses, two out of five mice in this group developed clear anemia indicated by the hematology panel. These findings suggest that excessive or poorly balanced activation of autophagy and mTOR inhibition can cross a toxicity threshold, potentially via impaired erythropoiesis or altered bone marrow function (Mitchell et al., 2016). Interestingly, aside from the adverse effects, the mice in this group did show some potentially positive signs, increased strength and social interaction. Rapamycin has been reported to reduce anxiety-like behavior in some studies, which may partially explain the observed increase in sociability (Chen et al., 2009; Halloran et al., 2012). However, any such benefit appears to be outweighed by the health risks observed at this dosage. Our protocol therefore highlights the importance of reformulating dose, route of administration, or treatment duration (e.g., intermittent dosing) before embarking on long-term survival studies.

Finally, pinealon, a short peptide with reported neuroprotective and nootropic properties (Arutjunyan et al., 2011), showed a modest trend toward improved working memory without obvious adverse effects. The absence of pronounced effects in this 8-week assay does not preclude meaningful benefits in older animals or under longer treatment durations. Instead, it highlights a key limitation of short-term performance screens: interventions that act primarily by delaying late-onset pathologies or reversing established frailty may require longer treatment windows or more advanced aging contexts to produce measurable functional improvements (Marcozzi et al., 2025; Mitchell et al., 2016).

In this line, several other limitations should be acknowledged. First, the use of only male C57BL/6J mice restricts conclusions about sex- and strain-specific responses, which are known to be substantial for many geroprotectors (Bartke et al., 2023; Knufinke et al., 2023). Second, the 8-week treatment window captures early and subchronic effects but may miss slower-developing benefits or toxicities, particularly in cognition and organ-specific pathology (Mitchell et al., 2016; Xu et al., 2021). Third, sample sizes were modest, reflecting the pilot nature of the screen; while sufficient to detect large and consistent phenotypes, but they limited statistical power to detect more modest or heterogeneous outcomes. Notably, lifespan and survival-related interpretations are highly sensitive to cohort variability and control performance, underscoring the importance of robust experimental design (Pabis et al., 2023). Fourth, phenotyping was intentionally limited, as we prioritized simple, cost-effective, and scalable readouts. However, additional measures such as body composition analysis, epigenetic clocks, inflammatory signatures, or organ-specific biomarkers could be incorporated depending on the nature of the intervention to strengthen mechanistic interpretation and translational relevance. Therefore, future iterations could include both sexes, increase sample size and treatment duration, and incorporate additional aging biomarkers tailored to the specific intervention under investigation.

An additional distinctive feature of this work is its funding model. To our knowledge, this is the first preclinical aging study in mice financed entirely through a tokenized decentralized science (DeSci) mechanism rather than conventional grant funding. DeSci frameworks use community-governed, on-chain funding structures to support research, aiming to allocate capital transparently and link contributors to scientific outcomes (Pump.Science and VitaDAO-like models)(Unfried, 2024). In the context of longevity research, where many projects are high-risk, high-reward and often underserved by conservative funding channels, this model offers a complementary pathway to support exploratory studies such as the present performance assay.

At a time when the longevity field is rapidly expanding and the number of putative anti-aging drugs, dietary paradigms, and genetic interventions continues to grow, this protocol addresses a major bottleneck in geroscience. It provides a scalable and practical pre-screening framework to prioritize the most promising candidates while minimizing translational and safety risks before initiating complex and resource-intensive lifespan trials. Importantly, this study also demonstrates how decentralized science infrastructures can move beyond conceptual models to support fully executed in vivo experimental pipelines conducted with strong scientific rigor.

Overall, compared to traditional mouse lifespan studies, which often require more than two years, costs exceeding $200K, and cohorts of over 100 animals, this approach reduced animal usage by 10-fold, shortened study duration to 4 months (6-fold faster), and decreased total costs to around $10K (20-fold less expensive). Together, these improvements provide a faster, and more economical and ethically sustainable pathway for early-stage longevity research.

## EXPERIMENTAL PROCEDURES

### Animal

All experiments were performed in male C57BL/6J mice obtained at 7 months of age (young adult, middle-age). Mice were group-housed (5 per cage) under standard specific-pathogen-free conditions (22 °C, 40–60% humidity, 12h/12h light-dark cycle) with ad libitum access to food and water. After the acclimation period (one month) and baseline testing, mice were randomly assigned (n = 5 per group) to either the Control group (standard diet, V1554, Ssniff) or one of five treatment groups. All interventions were initiated at the same age (8 months) and administered chronically for 8 weeks.

*EST: 17α-Estradiol* (IN-DA003NS5, Indagoo) – provided via food at 22 ppm concentration (approximately 2.5 mg/kg/day dose).

*R+SM: Rapamycin (encapsulated) + Smer28* (16-0202-027-P4, Rapamycin Holding and Indagoo, IN-DA003168) provided via food (rapamycin 22 ppm ≈ 2.5 mg/kg/day, plus Smer28 88 ppm ≈ 10 mg/kg/day).

*B+RE: Berberine + Resveratrol* (54-OR1029863 and 54-BIK9013 Apollo Scientific), provided via food, resveratrol 170 ppm ≈ 20 mg/kg/day; berberine 375 ppm ≈ 43 mg/kg/day.

*SIL: Sildenafil* (1130125, Acofarma) provided via food at 62 ppm ≈7 mg/kg/day.

*PNL: Pinealon (Glu-Asp-Arg)* (SKU:500, Chemiax) administered by intraperitoneal (i.p.) injection at 30 mg/kg/day. Pinealon was injected twice per week (Monday and Thursday), alternating between the left and right sides.

For dietary treatments, compounds were dissolved in a 1:1 water:soybean-oil (R111S041, Ssniff) solution and thoroughly mixed with the chow diet. Control and pinealon-treated animals received chow prepared with the same vehicle solution.

All animal experiments were conducted in accordance with institutional and national guidelines. Specifically, the study was carried out in Santiago de Compostela, Spain, under the approval of the competent authority (Expediente no. 15012/2025/017, Xunta de Galicia). The procedures complied with the European Union Directive 2010/63/EU on the protection of animals used for scientific purposes, as transposed into Spanish law by Royal Decree 53/2013.

#### Food and Water Intake

Mice were housed five per cage; therefore, food and water consumption were measured at the cage level. Food intake was calculated biweekly and normalized per animal (grams per day per mouse). Water intake (mL per day per mouse) was measured in a similar manner. Due to cage-level measurements and potential spillage, water intake data showed higher variability and were interpreted qualitatively, whereas food consumption measurements were considered reliable.

### Physiological Measurements

#### Body Weight and Temperature

Body weight (BW) was measured every other week at the same morning hour. Body temperature (BT) was measured at the same time points using a rectal thermometer for mice (BIO-TK8851 & BIO-BRET-3, Bioseb). Baseline BW and BT were recorded for 8 weeks prior to treatment and then monitored throughout the 8-week treatment period. For analysis, changes in BW and BT were calculated for each animal relative to its pretreatment baseline values.

### Behavioral and Functional Tests

We performed a battery of **functional assays** to evaluate physical, cognitive, and behavioral performance. All tests were conducted during the light phase. Baseline measurements were obtained just prior to treatment onset (Week 0), and end-point measurements were collected during the final week of treatment (Week 8) (Fig. 1a).

#### Open Field Test (Locomotor Activity)

Spontaneous activity was measured in an open-field arena (45 cm square) for 10 min. Total distance traveled (meters) and maximum speed (meters/seconds) achieved were quantified by an automated tracking system (Anymaze Tracking System, Stoelting). This test was performed to detect changes in general locomotor activity and endurance.

#### Grip Strength (Muscular Strength)

Forelimb grip strength was measured using a digital grip strength meter (BIO-GS4, BIO-GRIPGS, Bioseb). Each mouse was allowed to grasp a metal grid and was gently pulled backward until the grip was released; the peak force (g) was recorded. Two pulls were performed per session, and the maximum value was used for analysis. Grip strength was assessed repeatedly during the 8-week pretreatment period to establish baseline values and again during the treatment phase. For analysis, grip strength was normalized to body weight and expressed as percentage change relative to baseline.

#### Y-Maze (Cognitive Function)

Spatial working memory was assessed using the spontaneous alternation Y-maze test (three arms positioned at 120°). Each mouse was placed at the center of the maze and allowed to explore freely for 5 min. The sequence of arm entries was recorded. Working memory performance was quantified as the percentage of spontaneous alternations, defined as the proportion of overlapping triplets of consecutive arm entries that included all three arms without repetition. The total number of arm entries was also measured as an index of locomotor activity during the task. Y-maze testing was performed at week 8 only (no baseline measurement), as cognitive impairments were not expected to be pronounced at this age, and the assay was primarily intended to detect potential treatment-induced improvements.

#### Social Recognition Test

Social motivation and sexual preference were assessed in a square open-field arena (45cm x 45cm). Four stainless-steel baskets (9.5 cm diameter x 20 height) were place in the arena: two baskets on the left and right sides each contained an unfamiliar conspecific (male on the left, female on the right), and two identical baskets (top and bottom positions) were with an inanimate object. The test mouse was placed in the center and allowed to explore for 7 min. Body temperature was measured immediately before and after the test to assess thermoregulatory responses. Behavior was recorded using a video-tracking system for automated data collection (Anymaze Tracking System, Stoelting). Time spent exploring each stimulus (male, female and object baskets) was quantified; exploration was defined as sniffing, touching or orienting the nose toward the basket within 3 cm. No dedicated habituation session was performed; however, mice had been exposed to the same testing arena during an open-field session conducted a few days prior to testing. Social testing was done after 7 weeks of chronic treatment.

#### Blood Collection and Analysis

Baseline blood samples were collected prior to treatment initiation via submandibular vein puncture (cheek sampling) using a 25G needle. Approximately 150 µL of blood was collected into Microvette® 500 EDTA tubes under gentle restraint. At the endpoint (after 8 weeks of treatment), mice were euthanized by CO_2_ inhalation followed by decapitation. In both cases, mice were fasted for 4 hours prior to blood collection. Complete blood count (CBC) measurements were performed using an automated veterinary hematology analyzer (BC-5000 Vet, Mindray). The following parameters were obtained: white blood cell count (WBC, 10^9^/L), lymphocytes (Lym, 10^9^/L), monocytes (Mon, 10^9^/L), eosinophils (Eos, 10^9^/L), basophils (Bas, 10^9^/L), red blood cell count (RBC, 10^12^/L), hemoglobin (HGB, g/L), hematocrit (HCT, %), mean corpuscular volume (MCV, fL), mean corpuscular hemoglobin (MCH, pg), mean corpuscular hemoglobin concentration (MCHC, g/L), red cell distribution width (RDW-CV, %; RDW-SD, fL), platelet count (PLT, 10^9^/L), mean platelet volume (MPV, fL), and platelet distribution width (PDW, fL).

Following CBC analysis, blood samples were centrifuged at 6,500 rpm for 5 minutes to separate plasma, which was stored at −80 °C until further analysis. Fasting plasma glucose was measured using a glucose oxidase method (Sinocare Safe AQ Smart glucometer and test strips).

#### Urinalysis

Urine samples were collected immediately prior to euthanasia by gently restraining each mouse and collecting spontaneously voided urine. Because not all mice urinated and sample volumes varied, urine was pooled by treatment group. Urine was analyzed using an automated urine analyzer with corresponding urine test strips (VETSCAN UA, Abaxis). The following parameters were obtained: leukocytes (LEU, cells/μL), ketones (KET, mg/dL), nitrites (NIT), urobilinogen (URO), bilirubin (BIL, mg/dL), glucose (GLU, mg/dL), protein (PRO, mg/dL), specific gravity (SG), pH, blood (BLD, cells/μL), ascorbic acid (ASC, mg/dL), microalbumin (MA, mg/dL), calcium (Ca, mg/dL), creatinine (CR, mg/dL), and the protein/creatinine ratio (PRO/CR).

#### Mouse imaging

Mice were placed on a flat, well-lit surface and photographed using a tablet camera (Xiaomi Pad 7). During image acquisition, mice were gently restrained by the tail as needed to minimize movement and ensure consistent positioning. Pictures were obtained from multiple angles for documentation.

### Statistical Analysis

Data are presented as mean ± standard error of the mean (SEM). For each dataset, normality was assessed using the Shapiro-Wilk test. For normally distributed outcomes, homogeneity of variances was evaluated, and group differences were tested using one-way ANOVA (equal variances) or Welch’s ANOVA (unequal variances), followed by Dunnett’s post hoc test for comparisons versus the control group. For non-normally distributed outcomes, group differences were assessed using the Kruskal-Wallis test followed by Dunn’s post-hoc test for comparisons versus the control group. For repeated-measures outcomes, treatment and time effects were analyzed using two-way ANOVA when complete data were available; when missing values were present, a mixed-effects model (REML) was used. A significance threshold of *P* < 0.05 was used. Given the small sample size (*n*=5 per group), our analysis focused on large effect sizes and consistent trends.

## FIGURE LEGENDS

The legends for all main and supplementary figures and tables are included in their respective files.

## AUTHOR CONTRIBUTIONS

E.M.-J. performed all experiments, collected samples and data, and conducted statistical analyses. J.R.-C. and A.P. analyzed the data and wrote the manuscript with intellectual input from all authors. F.M.-R. organized and managed the project. P.P.-B. contributed to figure preparation and visual design. B.L. supported the experimental design and interpretation of results. A.P. conceived and supervised the study and prepared the final figures.

## ACKNOWLEDGMENTS

We thank the staff of the Mouse Facilities at the Centro de Biomedicina Experimental de la Universidad de Santiago de Compostela (CEBEGA) for enabling the use of their facilities, housing the mice, and assisting with sample analysis and equipment use.

