## Extended Data for "Short-Term Performance Assay Identifies Functional Benefits and Early Toxicity of Longevity Interventions in Mice"

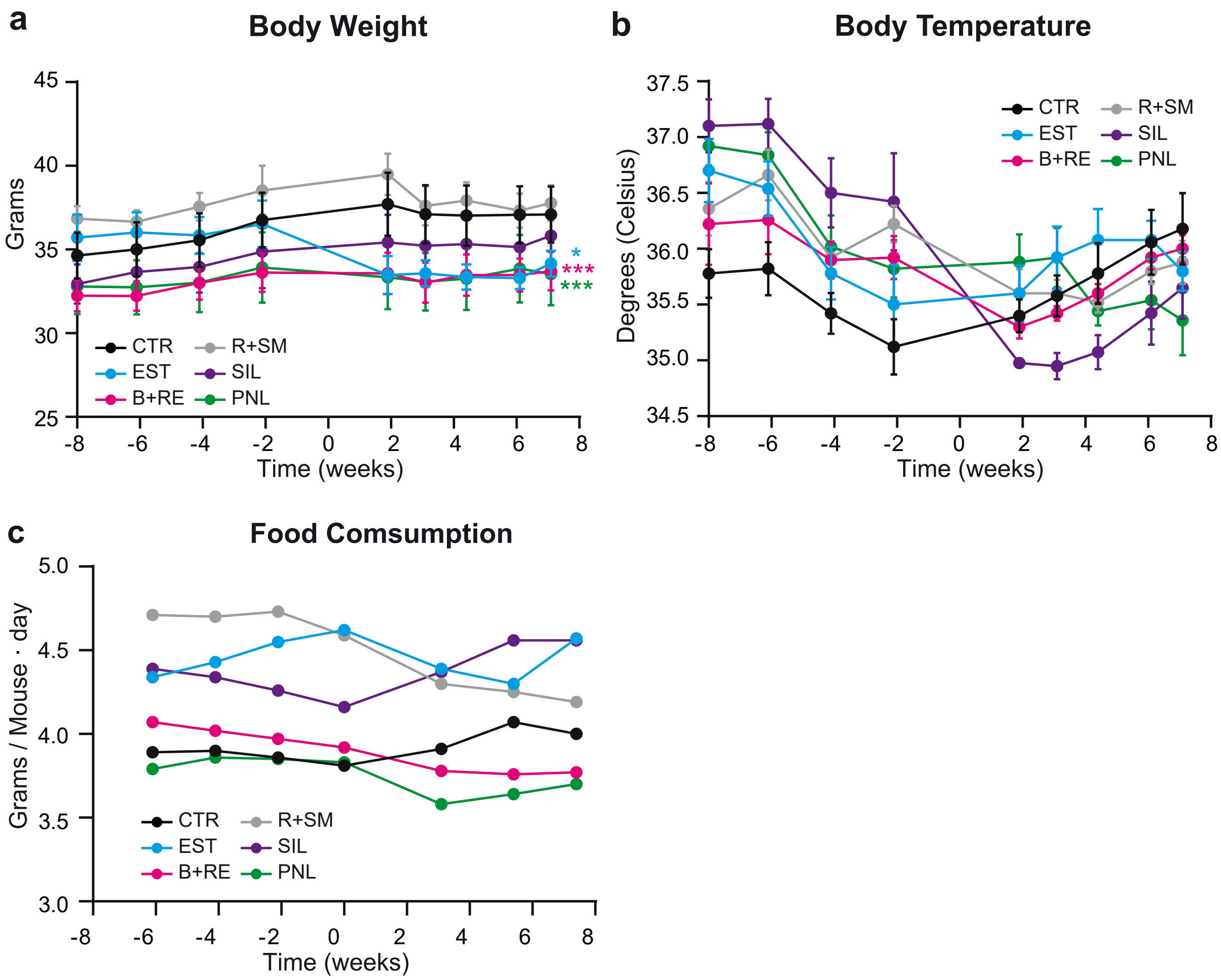


**Extended Data Figure 1 | Supplemental Basic physiological parameters. a,** Body weight in grams (g) measured and **b,** Rectal body temperature (°C) measured longitudinally throughout the protocol. **c,** Daily food intake (grams/mouse/day). CTR, control; EST, 17α-estradiol; B+RE, berberine + resveratrol; R+SM, rapamycin + Smer28; SIL, sildenafil; PNL, pinealon. Data show mean ± standard error of the mean (SEM). Statistical significance was assessed by mixed-effects model (REML) with Dunnett's multiple comparisons test **(a–b).**

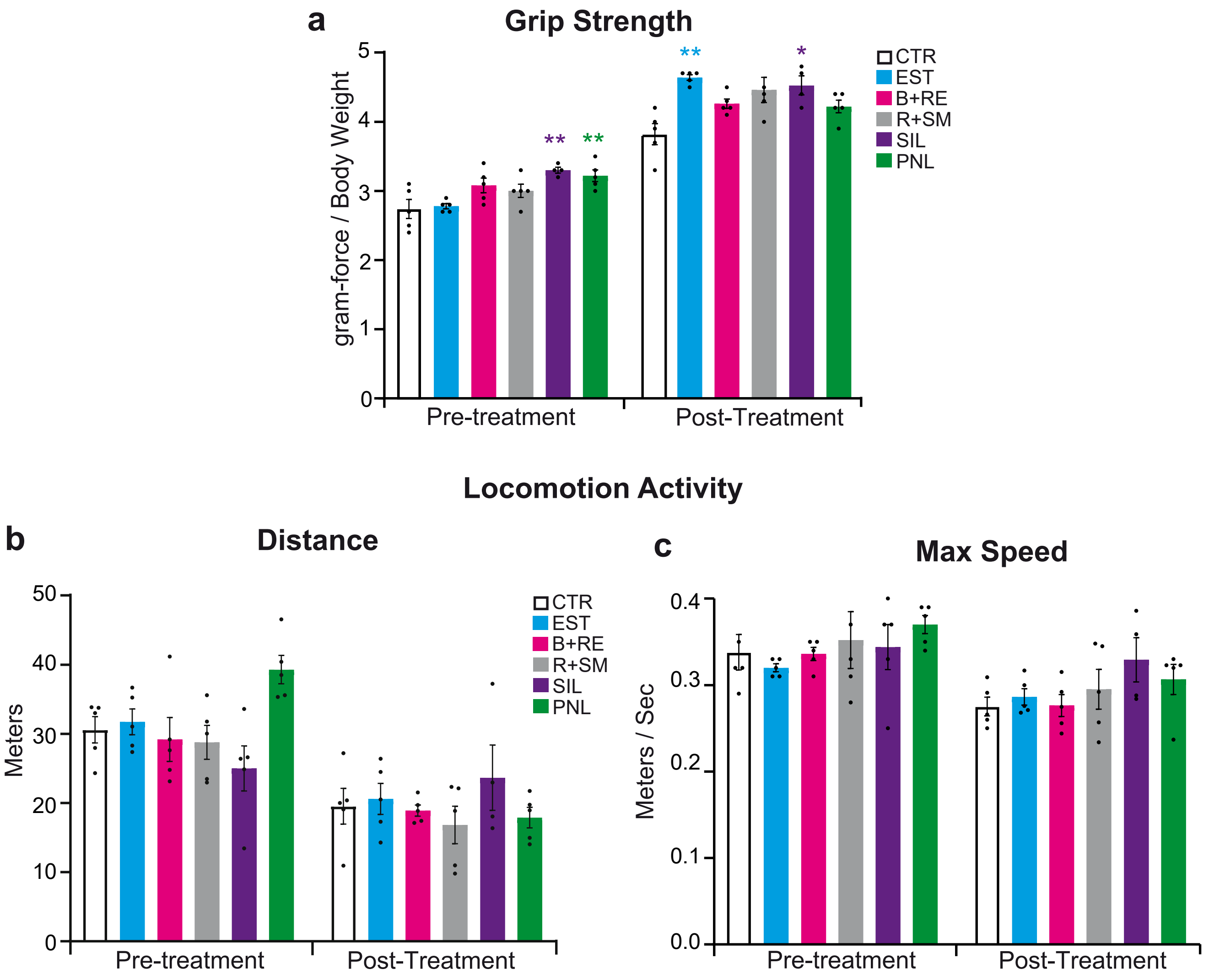


**Extended Data Figure 2 | Supplemental physical performance. a,** Forelimb grip strength normalized to body weight measured before (Pre) and after (Post) treatment. **b,** Total distance traveled and **c,** Maximum speed in the open field test measured pre- and post-treatment. CTR, control; EST, 17α-estradiol; B+RE, berberine + resveratrol; R+SM, rapamycin + Smer28; SIL, sildenafil; PNL, pinealon. Data show mean ± standard error of the mean (SEM). Statistical significance was assessed by one-way ANOVA with Dunnett's multiple comparisons test (**a-pre, b**), Kruskal-Wallis test with Dunn's multiple comparisons test (**a-post, c-post**) and Welch ANOVA with Dunnett's multiple comparisons test (**c-pre**).

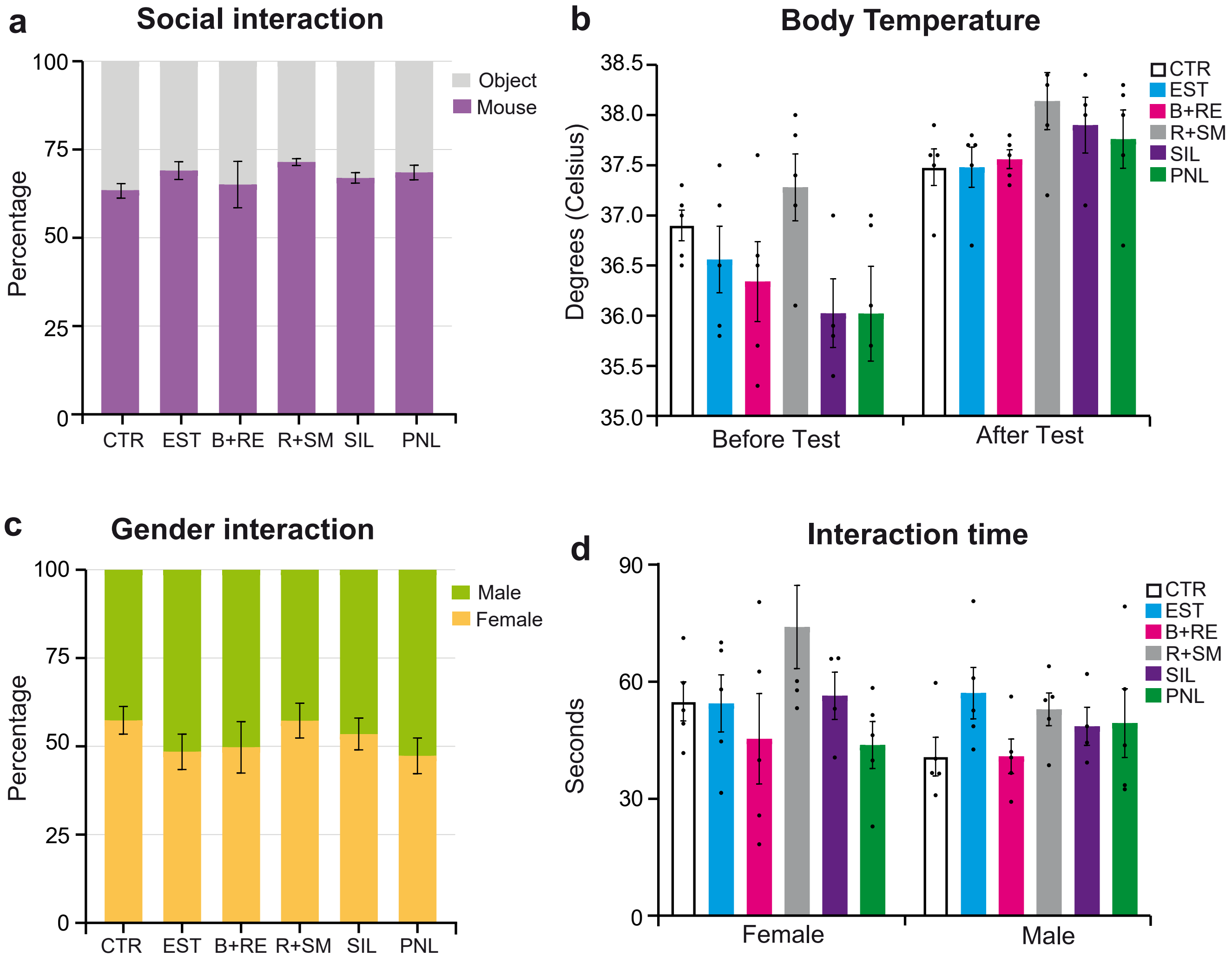


**Extended Data Figure 3 | Supplemental cognitive function and social behavior. a,** Percentage of interaction time with stranger mice versus objects during the social recognition test. **b,** Interaction time and **c,** Percentage of interaction time with female versus male stranger mice. **d,** Core body temperature measured before (Pre) and after (Post) the social recognition activity test. CTR, control; EST, 17α-estradiol; B+RE, berberine + resveratrol; R+SM, rapamycin + Smer28; SIL, sildenafil; PNL, pinealon. Data show mean ± standard error of the mean (SEM). Statistical significance was assessed by Kruskal-Wallis test with Dunn's multiple comparisons test (**a, b-after**) and one-way ANOVA with Dunnett's multiple test (**b-before, c, d**).

### Complete Blood Count

#### Pre-treatment (Baseline)

| Mouse ID | Group | WBC $10^9$/L | RBC $10^{12}$/L | HGB g/L | HCT (%) | MCV (fL) | RDW-CV (%) | RDW-SD (fL) | PLT $10^9$/L | MPV (fL) | PDW (fL) |
|---|---|---|---|---|---|---|---|---|---|---|---|
| **AVERAGE** | **CTR** | 8.726 | 10.082 | 148 | 43.84 | 43.54 | 18.38 | 38.6 | 1185.6 | 6.54 | 15.54 |
| **MOUSE_016** | R+SM | 10.94 | 10.68 | 153 | 45.6 | 42.7 | 18.7 | 38.8 | 1108 | 6.5 | 15.5 |
| **MOUSE_017†** | R+SM | 9.13 | **10.07** | **158** | **45.6** | 45.3 | 18 | 39.2 | 791 | 7.1 | 15.6 |
| **MOUSE_018** | R+SM | 11.25 | 10.63 | 152 | 45.5 | 42.8 | 17.5 | 36.3 | 1149 | 6.5 | 15.3 |
| **MOUSE_019†** | R+SM | 9.04 | **9.88** | **154** | **44.7** | 45.2 | 17.3 | 36.5 | 799 | 6.8 | 15.4 |
| **MOUSE_020** | R+SM | 11.91 | 10.95 | 156 | 46.7 | 42.7 | 18.3 | 38 | 1182 | 6.7 | 15.5 |

#### Post-treatment (Endpoint)

| Mouse ID | Group | WBC $10^9$/L | RBC $10^{12}$/L | HGB g/L | HCT (%) | MCV (fL) | RDW-CV (%) | RDW-SD (fL) | PLT $10^9$/L | MPV (fL) | PDW (fL) |
|---|---|---|---|---|---|---|---|---|---|---|---|
| **AVERAGE** | CTR | 8.032 | 9.63 | 142.75 | 41.925 | 43.98 | 16.56 | 34.34 | 579.5 | 6.6 | 16.08 |
| **MOUSE_016** | R+SM | 7.08 | 8.62 | 134 | 36.7 | 42.6 | 17 | 34.5 | 263 | 6.6 | 16.5 |
| **MOUSE_017†** | R+SM | 4.98 | **5.94** | **94** | **27.6** | 46.5 | 17.7 | 38.4 | 472 | 6.9 | 15.7 |
| **MOUSE_018** | R+SM | 11.17 | 9.43 | 138 | 40 | 42.4 | 19.2 | 38.6 | 520 | 6.4 | 15.3 |
| **MOUSE_019†** | R+SM | 7.71 | **6.54** | **116** | **31** | 47.4 | 17.9 | 39.6 | 375 | 7.2 | 15.6 |
| **MOUSE_020** | R+SM | 10.44 | 10.23 | 147 | 44.3 | 43.3 | 17.9 | 37 | 768 | 6.7 | 16.2 |

**Supplementary Table 1 | Complete blood count analysis of rapamycin + SMER28–treated mice.** Complete blood count (CBC) parameters measured at baseline (Pre-treatment) and after 8 weeks of treatment (Post-treatment) in rapamycin + Smer28 (R+SM) treated mice. Control values correspond to the mean of the control group (CTR). Mice showing hematological values consistent with anemia at endpoint are marked with † and highlighted in red. WBC, white blood cells; RBC, red blood cells; HGB, hemoglobin; HCT, hematocrit; MCV, mean corpuscular volume; RDW-CV, red cell distribution width; RDW-SD, red cell distribution width; PLT, platelets; MPV, mean platelet volume; PDW, platelet distribution width.

### Urinalysis parameters

Post-treatment (Endpoint)

| Group | LEU (cell/μ | KET (mg/dL | NIT | URO | BIL (mg/dL | GLU (mg/dL | PRO (mg/dL | SG | pH | BLD (cell/μ | ASC (mg/dL | MA (mg/dL | Ca (mg/dL | CR (mg/dL |
|---|---|---|---|---|---|---|---|---|---|---|---|---|---|---|
| EST | 0 | 0 | 0 | Normal | 0 | 0 | 15 | 1.01 | 6.5 | 0 | 0 | <2.5 | 30 | 50 |
| B+RE | 0 | 0 | 0 | Normal | 0 | 0 | <15 | 1.02 | 6 | 0 | 10 | <2.5 | 30 | 50 |
| R+SM | 0 | 0 | 0 | Normal | 0 | 0 | <15 | 1.02 | 5.5 | 0 | 0 | <2.5 | 20 | 50 |
| SIL | 0 | 0 | 0 | Normal | 0 | 0 | <15 | 1.015 | 6 | 0 | 0 | <2.5 | 20 | 50 |
| PNL | 0 | 0 | 0 | Normal | 0 | 0 | <15 | 1.02 | 6.5 | 0 | 0 | <2.5 | 30 | 50 |

**Supplementary Table 2 | Urinalysis parameters.** Urine analysis was performed after 8 weeks of treatment at the end point. Samples correspond to pooled urine collected from animals within the same treatment group. EST, 17α-estradiol; B+RE, berberine + resveratrol; R+SM, rapamycin + Smer28; SIL, sildenafil; PNL, pinealon. Parameters include: LEU, leukocytes; KET, ketones; NIT, nitrites; URO, urobilinogen; BIL, bilirubin; GLU, glucose; PRO, protein; SG, specific gravity; BLD, blood; ASC, ascorbic acid; MA, microalbumin; Ca, calcium; CR, creatinine.
